# FIRM: Flexible Integration of single-cell RNA-sequencing data for large-scale Multi-tissue cell atlas datasets

**DOI:** 10.1101/2020.06.02.129031

**Authors:** Jingsi Ming, Zhixiang Lin, Jia Zhao, Xiang Wan, Can Yang, Angela Ruohao Wu

## Abstract

Single-cell RNA-sequencing (scRNA-seq) is being used extensively to measure the mRNA expression of individual cells from deconstructed tissues, organs, and even entire organisms to generate cell atlas references, leading to discoveries of novel cell types and deeper insight into biological trajectories. These massive datasets are usually collected from many samples using different scRNA-seq technology platforms, including the popular SMART-Seq2 (SS2) and 10X platforms. Inherent heterogeneities between platforms, tissues, and other batch effects makes scRNA-seq data difficult to compare and integrate, especially in large-scale cell atlas efforts; yet, accurate integration is essential for gaining deeper insights into cell biology. Through comprehensive data exploration, we found that accurate integration is often hampered by differences in cell-type compositions. Herein we describe FIRM, an algorithm that addresses this problem and achieves efficient and accurate integration of heterogeneous scRNA-seq datasets across multiple tissue types, platforms, and experimental batches. We applied FIRM to numerous large-scale scRNA-seq datasets from mouse, mouse lemur, and human, comparing its performance in dataset integration with other state-of-the-art methods. FIRM-integrated datasets show accurate mixing of shared cell type identities and superior preservation of original structure without overcorrection, generating robust integrated datasets for downstream exploration and analysis. It is also a facile way to transfer cell type labels and annotations from one dataset to another, making it a reliable and versatile tool for scRNA-seq analysis, especially for cell atlas data integration.

---

The advent of single-cell RNA-sequencing (scRNA-seq) technology has enabled discovery of new cell types^1^, understanding of dynamic biological processes^2,3^ and spatial reconstruction of tissues^4^. Ongoing advancement in scRNA-seq technology has led to vast improvements in the scale and cost of the experiments^5–8^, providing unprecedented opportunities for deep biological insight. As scRNA-seq becomes more widely accessible both in availability and cost of the technology, many single cell transcriptomic datasets have now been generated for the same tissue types in various organisms, but often using different techniques and technology platforms. Prominent examples of this are recent efforts to generate cell atlases for whole organisms, including human^9–11^, mouse^12–15^ and mouse lemur. These cell atlas projects have generated scRNA-seq datasets encompassing a comprehensive set of tissues from the organism of interest, typically using many samples over multiple experiments. To ensure both technical sensitivity and scale in cell numbers profiled, many of these atlas projects, such as Tabula Muris^12^, Tabula Muris Senis^15^ and Tabula Microcebus consortium projects, also employ multiple different single-cell profiling technology platforms, including SMART-seq2 (SS2) and 10X Chromium (10X). Integrating datasets from different tissue types, samples and experiments, and from different platforms, not only enables the transfer of information from one dataset to another, such as the transfer of cell-type labels and annotations, but also makes the atlases more comprehensive and cohesive, which then benefits downstream biological analyses. However, complex technical variations and heterogeneities that exist between datasets makes integration challenging.

Existing methods have been designed for integration of scRNA-seq datasets across different samples, experiments, species or types of measurement. For example, batch correction methods based on mutual nearest neighbor in the original high-dimensional space, such as MNN^16^ and Scanorama^17^; methods by identification of shared lowdimensional space, such as CCA^18^, ZINB-WaVE^19^ and scVI^20^; methods which combine the previous two approaches to identify shared subpopulations in the low-dimensional space, such as Seurat^21^ and LIGER^22^; Harmony^23^ which corrects the low-dimensional embedding of cells; BBKNN^24^ which performs batch correction at the neighborhood graph step; and BUSseq^25^ which is based on Bayesian hierarchical models^23^. Although all these methods are applicable, they do not account for integration of datasets across multiple platforms. Specifically, SS2 and 10X are two frequently used scRNA-seq platforms with their unique strengths and weaknesses. SS2 is a plate-based full-length approach with high transcriptome coverage per cell and greater sensitivity^26^, whereas the microfluidic droplet-based method, 10X, generally has lower coverage per cell and a higher dropout rate^27^. But 10X is able to profile hundreds of thousands of cells per study with low per cell costs^8^, which enables more reliable detection of rare cell types, and the inclusion of unique molecular identifiers (UMIs) in 10X allows removal of amplification bias and in turn enables more accurate transcript abundance quantification^28^. Harmonizing datasets across multiple platforms for integrative analysis can take advantage of the strengths of each technology and improve the robustness, as well as achieve higher accuracy for visualization; better comparison across datasets and studies; and higher statistical power for differentially expression analysis. Furthermore, integration would make it possible to use 10X for discovery of new cell types, while taking advantage of the greater depth and sensitivity of SS2 to investigate their biology, including enabling analysis of transcript isoforms, splicing^29–31^, and allelic expression^30,32^. This is particularly important for large-scale cell atlas projects, which are intended to serve as robust and comprehensive reference datasets for future mining. Due to technical variations and characteristic differences in SS2 and 10X datasets, not accounting for platform-specific characteristics during integration can lead to inaccuracies under different scenarios: sometimes resulting in poor alignment of cells from the same cell type; other times mixing cells from different cell types inappropriately, giving rise to overcorrection. An ideal method requires identification of the main technical variation for integration and designing a specific approach to address it.

Through comprehensive data exploration, we found that the differences in depth of expression profiles are the main technical variation between SS2 and 10X datasets and the heterogeneity in cell type composition accounts for the main problem preventing accurate integration. Datasets with different cell type compositions have different directions of maximum variance chosen by principal component analysis (PCA) and perform differently after standard preprocessing procedures including normalization and scaling. We have developed an efficient algorithm, FIRM, to specifically account for this composition effect thereby harmonizing SS2 and 10X datasets. Authors of other methods such as MNN and Scanorama have also observed the influence of cell type composition on integration and tried to reduce this effect by modifying the underlying expression data to align cells with high similarity. However, using this approach, other problems such as overcorrection can occur, especially when there are dataset-specific cell types. This is a common occurrence in cell atlas projects that require integration of different tissue types. In overcorrecting, close but not identical cell types may be merged into the same cluster inappropriately. In contrast, FIRM applies a re-scaling procedure based on subsampling for both datasets in a unified workflow. Overcorrection can be avoided with this approach and the original structure for each dataset can be largely preserved in the integrated dataset, generating a reliable input for downstream analysis. We applied FIRM to integrate numerous scRNA-seq datasets generated using different platforms and sample types. Compared with existing state-of-the-art methods, FIRM not only demonstrates great integration performance but also effectively avoid overcorrection in all tested datasets.

## Results

### Differences in cell type composition is a major factor preventing accurate integration of scRNA-seq data generated by different technology platforms

To specifically investigate the influence of cell type composition on scRNA-seq dataset integration outcomes, we consider a toy example with two scenarios using hypothetical datasets in which all technical variations have been removed except the difference in their cell type compositions. In the first scenario, the cell type proportions are consistent across different platforms (SS2: 50% cell type 1 + 50% cell type 2, 10x: 50% cell type 1 + 50% cell type 2); in the second scenario, the cell type proportions are different (SS2: 50% cell type 1 + 50% cell type 2, 10x: 80% cell type 1 + 20% cell type 2). We scaled the expression value for each gene to unit variance for each dataset, which is the standard preprocessing procedure applied to prevent the dominance of highly expressed genes and is also necessary to reduce the difference in sequencing depth for dataset integration across platforms. In the first scenario, where cell type compositions in SS2 and 10X datasets are the same, cells belonging to the same cell type have similar gene expression levels after scaling and are well mixed across platforms (Fig. 1a). This is the ideal case for downstream analysis, in which only the biological variations among cell types are reserved. However, in the second scenario, the scaled expression values in SS2 and 10X datasets for cells of the same type show large differences because of their heterogeneity in cell type composition, resulting in poor integration of these two datasets (Fig. 1b). This demonstrates that when cell-type composition is skewed between the two datasets being integrated, it impacts the integration outcome and can result in inaccurate cell merging.

**Fig. 1.**
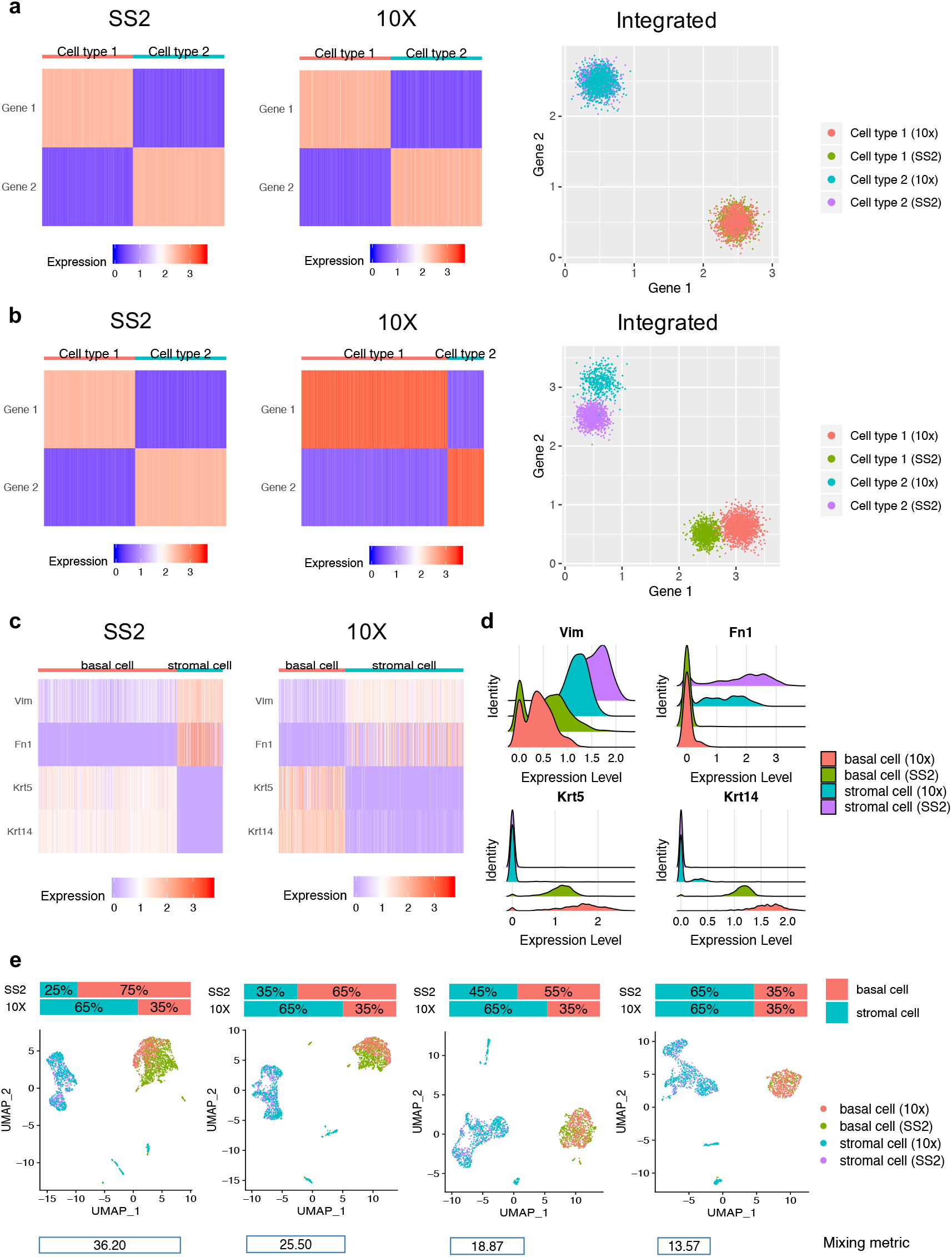
Illustration of the influence of cell type composition for scRNA-seq datasets integration based on hypothetical datasets (a-b) and real datasets (c-e). **a-b**, Gene expressions for cells in SS2 dataset, 10X dataset and integrated dataset after scaling to unit variance for each gene, when the cell type compositions in the hypothetical datasets are the same across datasets (SS2: 50% cell type 1 + 50% cell type 2; 10X: 50% cell type 1 + 50% cell type 2) (**a**) and when the cell type compositions are different across datasets (SS2: 50% cell type 1 + 50% cell type 2; 10X: 80% cell type 1 + 20% cell type 2) (**b**). **c-e**, Illustration of the key problem for integration based on the mammary gland scRNA-seq datasets generated by SS2 and 10X from Tabula Muris, withholding only the basal cells and stromal cells. **c**, Marker expressions for basal cells and stromal cells in SS2 dataset and 10X dataset after scaling to unit variance for each gene, where the cell type compositions are different across datasets (SS2: 75% basal cells + 25% stromal cells; 10X: 35% basal cells + 50% stromal cells). **d**, Ridge plots for marker expressions for each cell type in each dataset. **e**, Uniform manifold approximation and projection (UMAP) visualization and mixing metric for the integrated dataset with different cell type composition by subsampling basal cells in SS2 dataset.

To verify our hypothesis using real scRNA-seq datasets across multiple platforms, we analyzed the Tabula Muris^12^ mouse mammary gland scRNA-seq data that was generated using SS2 and 10X, in which the relative proportions of basal cells and stromal cells across platforms are vastly different (SS2: basal cells/stromal cells = 75%/25%; 10X: basal cells/stromal cells = 35%/65%). We extracted the basal cells and stromal cells in this dataset to create a simple example resembling the previously illustrated toy example. After preprocessing each dataset, we compared the gene expression of two marker genes for stromal cells (*Vim* and *Fn1*) and another two for basal cells (*Krt5* and *Krt14*) between the SS2 and 10X dataset. We found that the expression levels for the same cell type marker across platforms are different in expression modes and dispersions (Fig. 1c, d). We then integrated the dataset by concatenating the scaled SS2 and 10X expression data matrices, and in visualizing the outcome we found that in this cell type composition scenario (SS2: 75% basal cells + 25% stromal cells; 10X: 35% basal cells + 65% stromal cells), basal cells across platforms did not correctly merge into one single cluster (Fig. 1e, leftmost panel).

In order to confirm whether this poor alignment is caused by the difference in cell type proportions, we performed subsampling to gradually reduce the proportion of basal cells in SS2 dataset from 75% to 35%, to match that of the 10X dataset. Then we integrated the 10X dataset with these subsets of SS2 dataset, and evaluated the performance. In addition to the UMAP^33^ plot for visualization, we also calculated the mixing metric (see Methods) to measure how well the datasets mixed after integration, where a lower score typically indicates better mixing performance. We indeed observed that more consistent cell type proportions gave rise to better alignments (Fig. 1e). Therefore, we concluded that the effects of heterogeneity in cell type composition between SS2 and 10X datasets accounts for one of the main technical variation preventing accurate integration of scRNA-seq data across platforms.

### FIRM provides accurate mixing of shared cell type identities and preserves local structure for each dataset

FIRM harmonizes SS2 and 10X datasets while accounting for the difference in cell type composition. The alignment workflow takes two scRNA-seq expression matrices as the input, typically one SS2 and one 10X dataset, and performs the following steps: (i) For each dataset, we conduct the standard pre-processing procedure which includes normalization, scaling and feature selection; (ii) Then we perform dimension reduction for each dataset using PCA and cluster cells based on the obtained low-dimensional representations; (iii) In order to align clusters in 10X dataset with clusters in SS2 dataset representing the same cell types, we check the alignment via subsampling to avoid overcorrection; (iv) For each pair of aligned clusters, we subsample the cells within the cluster to ensure that cell-type proportions are the same in SS2 and 10X datasets, then based on these subsampled cells, we calculate the standard deviation to perform re-scaling on each of the full datasets; (v) Finally, we merge the scaled data to obtain the integrated dataset. More technical details are presented in the Methods section.

First, we examined the performance for the integration of SS2 and 10X scRNA-seq datasets generated from the same tissue type where most cell type identities are shared across platforms. We applied FIRM to numerous paired SS2 and 10X scRNA-seq datasets to demonstrate its superior performance in data integration compared with existing state-of-the-art methods, including Seurat^21^, Harmony^23^, BBKNN^24^, BUSseq^25^, LIGER^22^, Scanorama^17^, MNN^16^, scVI^20^ and ZINB-WaVE^19^. The datasets include 13 pairs of SS2 and 10X scRNA-seq datasets from Tabula Muris^12^, 25 pairs from Tabula Microcebus, and one pair in Human Lung Cell Atlas^34^ (Methods). To evaluate the performance of integration, in addition to the visualization using UMAP on the integrated or corrected data, we made use of four metrics (Methods): mixing metric, local structure metric, average silhouette width (ASW) and adjusted rand index (ARI). The mixing metric measures how well the datasets are merged after integration, where a lower score typically indicates better mixing performance. The local structure metric measures how well the original structure of each dataset was preserved after data integration, where a lower score indicates worse degree of preservation and higher probability of overcorrection. ASW and ARI are calculated based on cell identities given in the original papers^12,34^. Higher values of ASW indicate that cells of the same type are closer to each other and further apart from cells of other types. Higher values of ARI indicate higher similarities between the clustering of integrated data with the predefined cell types. As these metrics represent different aspects of performance, joint consideration is required for effective comparison. For example, there is a trade-off between the mixing metric and the local structure metric – a low mixing metric does not always mean accurate integration, since overcorrection is characterized by a low mixing metric and a low local structure metric.

For all datasets tested, FIRM outperforms or is comparable to all other benchmarked methods for integration of SS2 and 10X datasets with relatively low mixing metric, and high local structure metric, including ARI and ASW. (Fig. 2 and Supplementary Figs. 1-31; SI unpublished data is embargoed until publication). FIRM not only provides accurate mixing of shared cell type identities, but also achieves superior preservation of the local structure for each dataset, which is one of the greatest advantages of FIRM over other methods. This is because FIRM harmonizes datasets through a re-scaling procedure without smoothing the expression of similar cell types across datasets towards each other, so that the relative expression patterns across cells within each dataset can be largely preserved. For almost all (36 out of 39) the integrated datasets, we found that FIRM achieved the highest local structure metric compared with all other methods (Supplementary Fig. 31), indicating minimal distortion of the between-cell-type relationships within each dataset, thus providing more credible integrated data for downstream analysis. For other benchmarked methods, different situations arose indicating non-ideal integration, including poor mixing of shared cell types, inappropriate mixing of different cell types, and weak preservation of the original dataset structure.

**Fig. 2.**
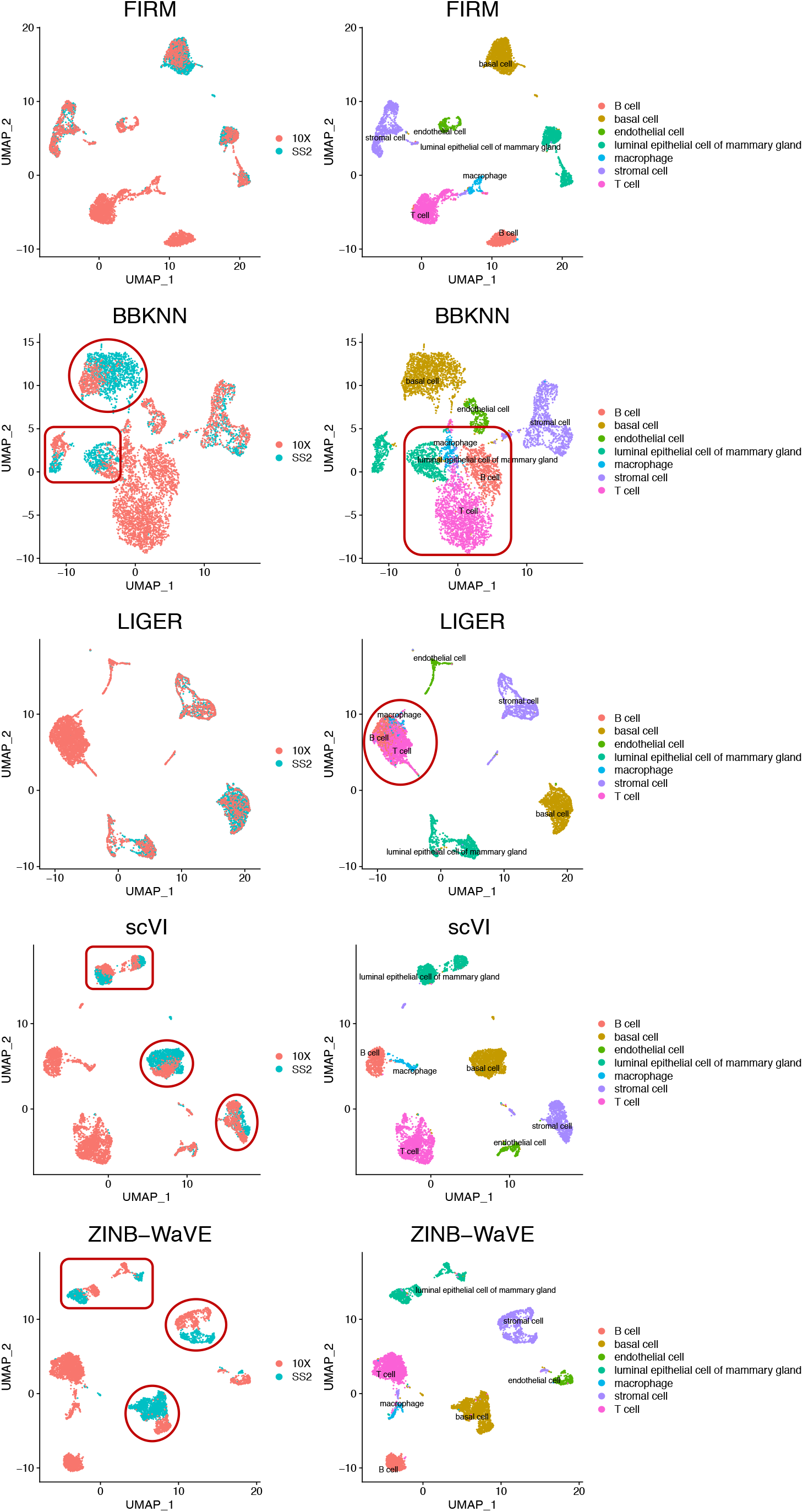
Comparison of integration methods based on the mammary gland scRNA-seq datasets generated by SS2 and 10X from Tabula Muris. UMAP plots of the integrated scRNA-seq dataset colored by platform (left panel) and by cell type (right panel) using FIRM, Seurat, BBKNN, BUSseq, LIGER, scVI and ZINB-WaVE.

Specifically, scVI and ZINB-WaVE are the two methods with the highest mixing metric, and this inadequate mixing of cell types can be seen in UMAP plots even by visual inspection (Fig. 2 and Supplementary Fig. 1-31). BBKNN is shown to have poor mixing performance when applying to the Tabula Muris bladder data (Supplementary Fig. 2), limb muscle data (Supplementary Fig. 3) and mammary gland data (Fig. 2); the Tabula Microcebus blood data and spleen data from lemur 2 (Supplementary Fig. 13, 17); and the Human Lung Atlas data (Supplementary Fig. 30). BUSseq also has relatively high mixing metric and worse mixing performance in many cases, including the Tabula Muris bladder data (Supplementary Fig. 2) and lung data (Supplementary Fig. 5); the Tabula Microcebus lung data and spleen data from lemur 2 (Supplementary Fig. 16, 17), eye retina data from lemur 4 (Supplementary Fig. 21); and the Human Lung Atlas data (Supplementary Fig. 30). Poor mixing performance was observed when applying Scanorama to the Tabula Muris spleen data (Supplementary Fig. 7), as well as to the Tabula Microcebus spleen data from lemur 2 and lemur 4 (Supplementary Fig. 17, 27).

LIGER overcorrected the datasets for some cases resulting in inappropriate mixing of different cell types, which is reflected by low ARIs. For example, LIGER incorrectly merged the B cells, macrophages, and T cells in the Tabula Muris mammary gland dataset (Fig. 2); the natural killer cells with the T-cells in the Human Lung Atlas (Supplementary Fig. 30); the subtypes of natural killer cells with the T-cells in the Tabula Microcebus kidney data and small intestine data from lemur 4 (Supplementary Fig. 23, 26); and the basal cells with the suprabasal cells in the Tabula Microcebus tongue data from lemur 4 (Supplementary Fig. 29). Harmony also has the phenomenon of overcorrection for some cases, for example, the inappropriate mixing of the cardiac muscle cells and erythrocytes in the Tabula Muris aorta dataset (Supplementary Fig. 1); the capillary cells and neutrophils in the Tabula Microcebus blood data from lemur 2 (Supplementary Fig. 13); the eosinophils and B cells in the Tabula Microcebus spleen data from lemur 4 (Supplementary Fig. 27).

For the preservation of original structure for each dataset, BUSseq was shown to have low local structure metrics (Supplementary Fig. 31), and is prone to separate the same type of cells, or different types of cells with a gradual transition, into discrete clusters. A few examples include the separation of the granulocytes and granulocytopoieic cells in the Tabula Muris marrow dataset (Supplementary Fig. 6); the mesenchymal cells in the Tabula Muris trachea dataset (Supplementary Fig. 8); the neutrophils in the Tabula Microcebus blood dataset from lemur 1 and lemur 2 (Supplementary Fig. 9, 13), and the bone marrow dataset from lemur 4 (Supplementary Fig. 20); the pachytene spermatocytes, round spermatids, elongating spermatids, and elongated spermatids in the Tabula Microcebus testes dataset from lemur 4 (Supplementary Fig. 28). MNN is prone to separate cells into small clusters and showed low ARIs for the blood, brain and lung data from lemur 1 (Supplementary Fig. 9-11); bladder data from lemur 2 (Supplementary Fig. 12); and pancreas data from lemur 4 (Supplementary Fig. 25) in Tabula Microcebus. Harmony suffers from the previous two problems in some cases. For example, Harmony discretizes the mesenchymal cells in the Tabula Muris trachea dataset (Supplementary Fig. 8), the neutrophils in the Tabula Microcebus blood and lung datasets from lemur 1, and bone marrow dataset from lemur 2 (Supplementary Fig. 9, 11, 14); Harmony also separates cells into small clusters for the heart and liver data from lemur 4 in Tabula Microcebus (Supplementary Fig. 22, 24). BBKNN is weak in separation of different cell types resulting in the lowest ARI in most cases (32 out of 39), including the mammary gland data in Tabula Muris (Fig. 2).

Harmony and Seurat are two popular methods with relatively better integration performance over other benchmarked methods. Compared with Harmony, FIRM shows its superiority in integration by achieving the lower mixing metric and higher local structure metric (Supplementary Fig. 32; SI embargoed until publication). Seurat is the method with the closest performance to FIRM (Supplementary Fig. 32). Seurat and FIRM have comparable performance in terms of ASW, but FIRM is superior in terms of ARI. Although Seurat usually has lower mixing metrics, FIRM does not show any obvious deficiency for mixing based on the UMAP plots of the integrated dataset. Considering the trade-off between the mixing metric and local structure metric, FIRM’s higher local structure metric suggests that it is more robust than Seurat in avoiding overcorrection.

### FIRM is robust against overcorrection

Cell atlas projects usually consist of scRNA-seq datasets for a comprehensive set of tissues, often spanning all the organs of an organism. The composition of cell types is largely different across tissues, while some cell types such as immune cells, fat cells, and cells of the vasculature are shared between multiple tissues or organs; cross comparison of different cell types and joint analysis of shared cell types across tissues are both valuable and informative. As such, it is essential that integration approaches not only accurately integrate datasets from multiple experiments or technology platforms, but also across different tissues.

Other integration methods, such as Seurat, LIGER and MNN, directly adjust the data matrices so that neighboring cells across different datasets have similar adjusted expression profiles, but this process of adjustment is vulnerable to overcorrection because the cells that are close in distance across datasets may not always be biologically similar. In the previous section, we found that for the same tissue when more than one cell type is shared across datasets, other benchmarked methods cannot completely avoid overcorrection. Different from other methods that project reference dataset onto query dataset based on neighboring cells across datasets, FIRM harmonizes datasets by incorporating scaling factors that account for differences in cell type compositions across datasets. As a result, FIRM can avoid overcorrection even if there are no shared cell types across the datasets being integrated, which is particularly important when integrating across multiple tissue types.

To evaluate whether the data integration methods are prone to overcorrection, we use the benchmark methods to integrate two datasets that had shared cell types manually removed, such that they have no cell types in common: SS2 dataset of kidney, and 10X dataset of brain cortex of lemur 2 in Tabula Microcebus. We applied FIRM, Seurat, Harmony, BBKNN, BUSseq, LIGER, Scanorama, and MNN to integrate these two datasets (Fig. 3); we excluded scVI and ZINB-WaVE from this assessment as these two methods did not work well even when there were shared cell types across datasets. Of all the methods assessed, FIRM and Harmony perfectly separated the cell types from each dataset and achieved high mixing metric. Other methods all suffered from overcorrection to varying degrees. Severe overcorrection was observed in Seurat, BUSseq, LIGER and MNN, where neurons and T-cells were incorrectly mixed. These other methods also inappropriately clustered different cell types together, resulting in low ARIs. The advantage of local structure preservation, one of the key strengths of the FIRM approach, is especially beneficial for integration across different tissues.

**Fig. 3.**
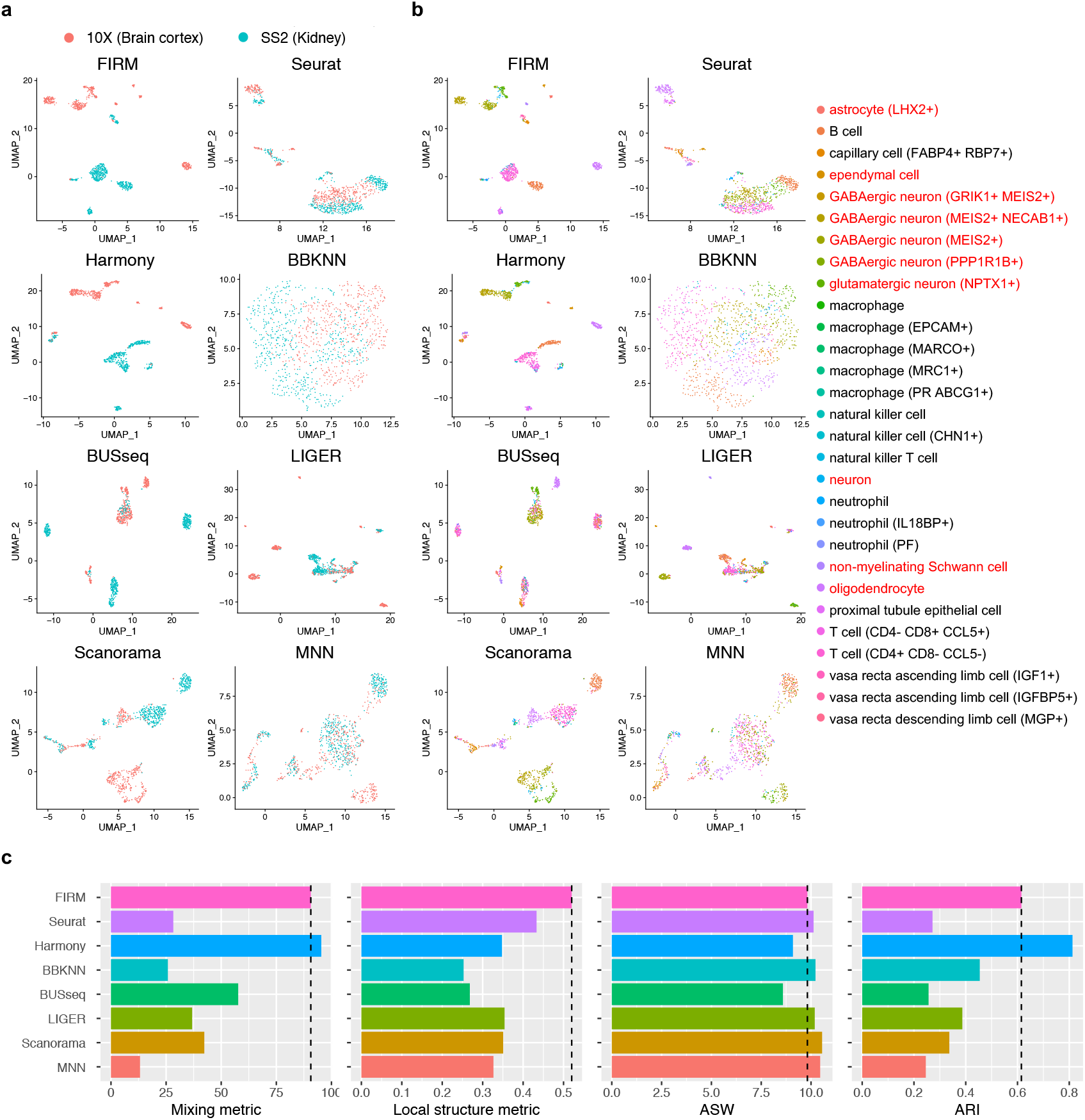
Comparison of integration methods for scRNA-seq datasets from two tissues in Tabula Microcebus (lemur 2) generated by different platforms (Kidney: SS2, Brain cortex: 10X. For clear illustration, we withheld several cell types in each of the dataset to make the cell types non-overlapped across datasets. **a**, **b**, UMAP plots of scRNA-seq datasets colored by platform (**a**) and by cell type (**b**) after integration using FIRM, Seurat, Harmony, BBKNN, BUSseq, LIGER, Scanorama and MNN. The labels for cell types in Brain cortex (10X) are colored by red. **c**, Metrics for evaluating performance across the six methods on four properties: cell mixing across platforms (Mixing metric), the preservation of within-dataset local structure (Local structure metric), average silhouette width of annotated subpopulations (ASW) and adjusted rand index (ARI). The dashed lines were set at the values for FIRM as reference lines.

### FIRM can transfer cell type identity labels across datasets and provide better clustering

By integrating SS2 and 10X datasets, we can take advantage of the strengths of each technology and improve data robustness. 10X datasets have higher throughput and usually more cell types are captured; in the SS2 dataset, some cell types may contain very few cells and fail to be identified if analyzed alone. Based on the SS2-10X integrated dataset, we can transfer information between datasets, such as cell type annotations and identity labels. One way to effectively label cell populations in SS2 data is by transferring the manually annotated 10X cell type identity labels to SS2 cells by detecting nearest neighbors for each SS2 cell in 10X dataset (Methods). For example, in the Tabula Microcebus testes SS2 dataset, we are not able to distinctly identify spermatogonia as there are only a few of them. By incorporating information from the 10X dataset, we identified three spermatogonia in the SS2 dataset that have marker expression patterns (*KIT, SOHLH1, PHOXF1, ZBTB16*) consistent with spermatogonia in the 10X dataset (Supplementary Fig. 33; SI embargoed until publication). We also noted that some marker genes (*OVOL1, SPO11, TEX101*) show clearer patterns in the SS2 dataset compared with the 10X dataset, indicating the benefit of detecting low abundance transcripts using SS2. For cases where the SS2 dataset contains more cell types than 10X dataset, we designed match scores such that cells with low scores can be labeled as ‘unknown’ (Methods).

FIRM can be applied to align more than two datasets, such as when harmonizing datasets generated from multiple individuals and platforms for one specific tissue. After accurate harmonization of multiple datasets, performing clustering on the integrated dataset can provide more reliable and consistent cluster labels for each dataset; taking advantage of the enhanced statistical power of the larger integrated dataset enables identification of rare cell types that may be missed in separated datasets. For example, kidney urothelial cells in the Tabula Microcebus are extremely rare in all the individual datasets: none from lemur 1, 4 cells from lemur 2, 4 cells from lemur 3, and 7 cells from lemur 4. They were not readily identifiable when the kidney datasets were individually annotated. After using FIRM to integrate all the kidney datasets across individuals and platforms, a small cluster of urothelial cells could then be detected with the specific markers (*UPK1A, FOXA1* and *UPK3A*) expressed (Fig. 4).

**Fig. 4.**
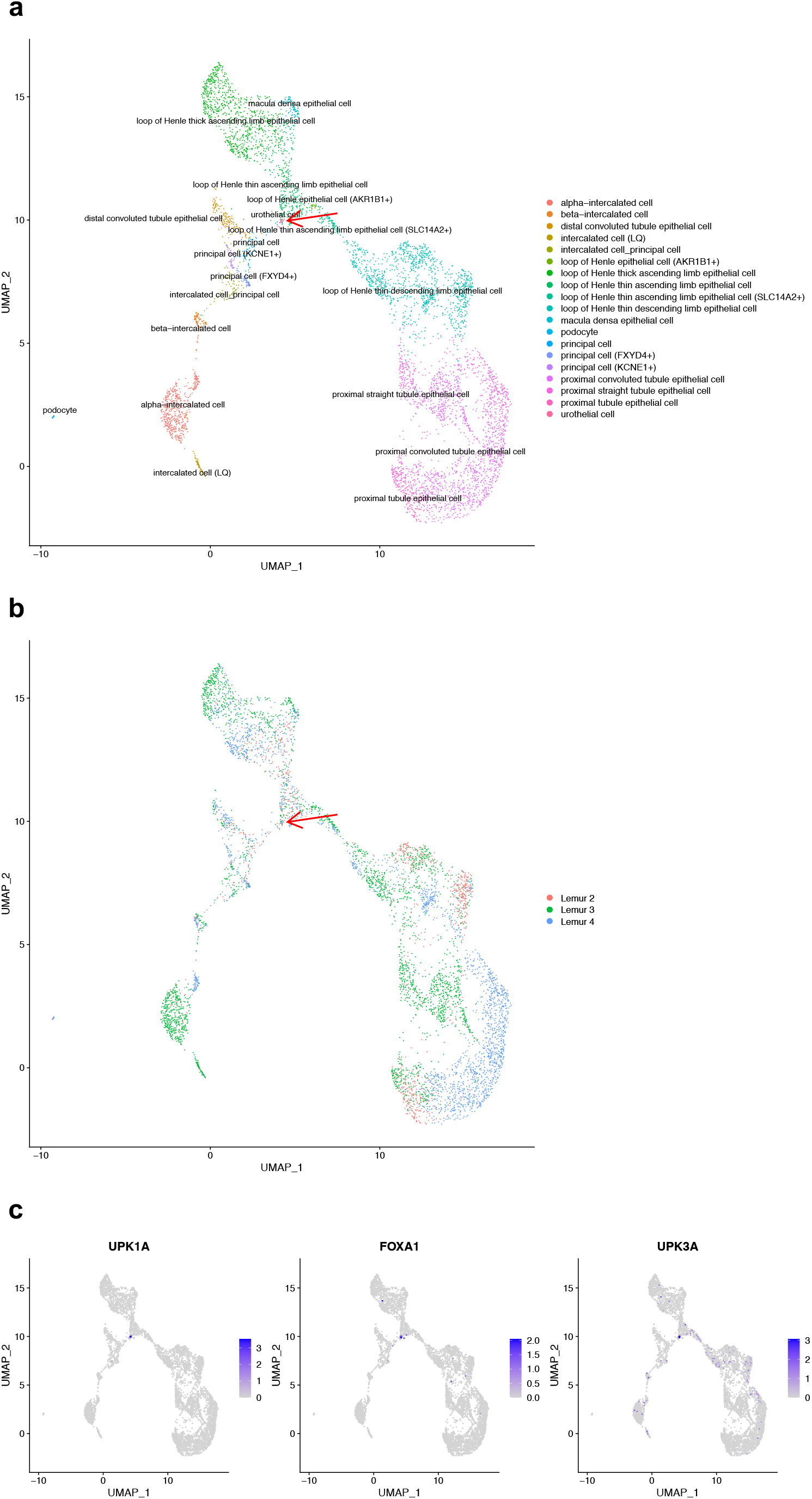
The FIRM integration for the kidney datasets across individuals and platforms in Tabula Microcebus. We subset the scRNA-seq datasets to keep the cells belonging to the epithelial compartment. **a**, **b**, UMAP plots colored by cell type (**a**) and by individual (**b**) after integration using FIRM. **c**, The expression levels of three marker genes (UPK1A, FOXA1 and UPK3A) for urothelial cells.

### FIRM accurately constructs cell atlases for entire organisms

FIRM can also be applied to harmonize datasets from multiple tissues in the cell atlas project to construct an atlas for the entire organism. We applied FIRM to integrate all the SS2 datasets from three individuals and 20 tissues in Tabula Microcebus and compared it with results from Seurat, Harmony, BBKNN and Scanorama, four popular methods for multiple datasets integration (Fig. 5). In this study, 29 SS2 datasets across individuals or tissues, which contain a total of 12,329 cells, were integrated. The integrated visualizations revealed that FIRM can provide accurate mixing of the shared cell types across both tissues and individuals, while preserving clear separation of various tissue compartments. For example, the germ cells which only exist in the testes dataset in this study can be viewed as a “sanity check”. FIRM separated the germ cells from other types of cells while retaining its structure from the original dataset. In contrast, Seurat suffered from severe overcorrection in merging cells from different compartments. Overcorrection also occurred when applying Harmony, for example, a few stromal cells were mixed in germ cells, epithelial cells were mixed in megakaryocyte-erythroid cells, hematopoietic cells were mixed in endothelial cells, myeloid cells were mixed in lymphoid cells. For BBKNN, the tissue compartments were close to each other, including the germ cells. Although Scanorama did not merge germ cells with other cells, the endothelial cells, epithelial cells, and stromal cells could not be distinguished from one another in the integrated result. Furthermore, Seurat may face difficulties integrating multiple large datasets with a very small dataset, such as those with less than 100 cells, because the numbers of neighbors selected for finding anchors are the same across datasets. For small dataset, only a small number of neighbors can be chosen, but this greatly influences the effectiveness when integrating other large datasets. Harmony only provides the integrated low-dimensional cell embeddings, rather than the expression profiles for the whole gene set, limiting its applicability for downstream analysis that require full gene expression profiles.

**Fig. 5.**
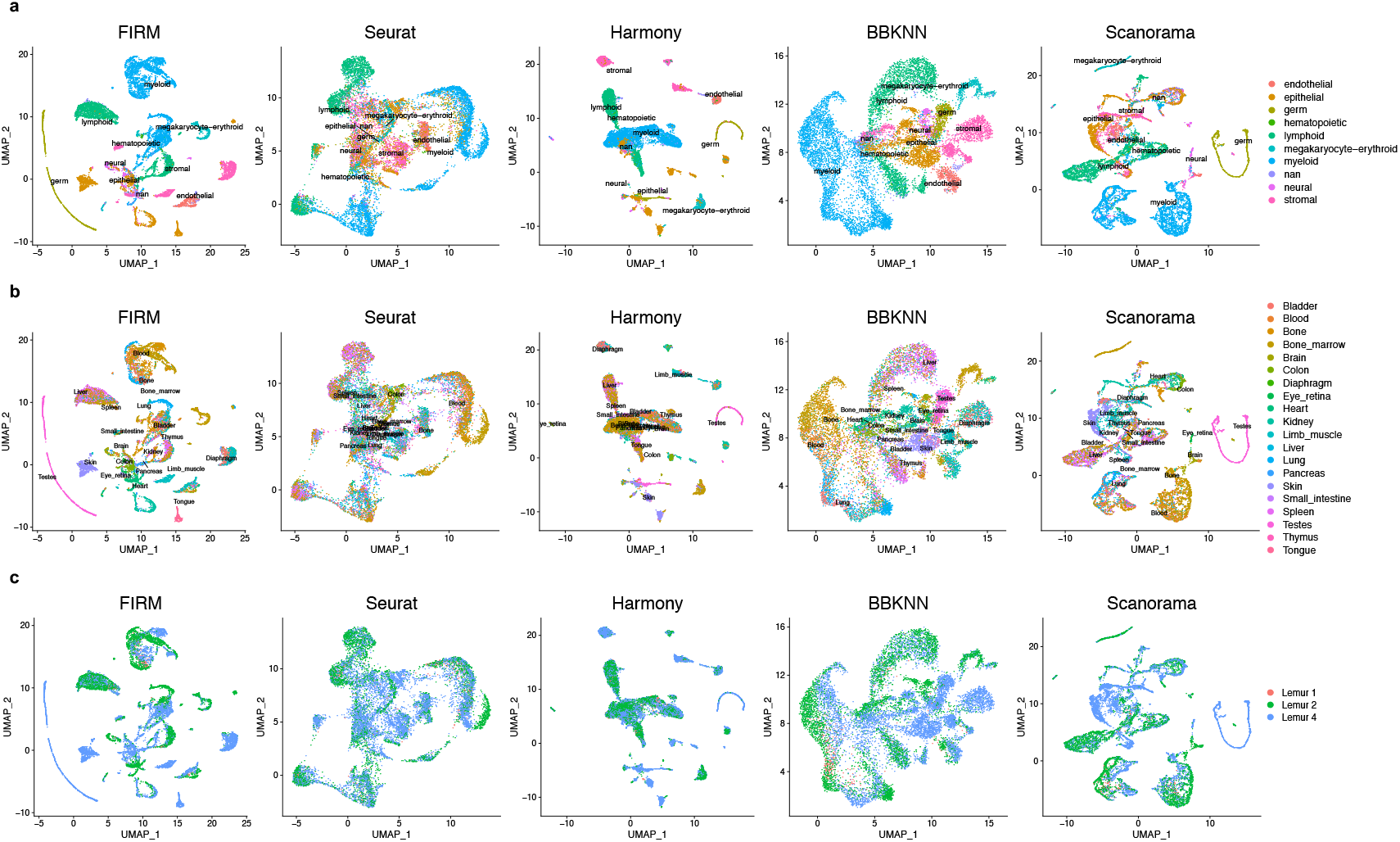
Comparison of FIRM, Seurat, Harmony, BBKNN and Scanorama for integration of all SS2 datasets across individuals and tissues in Tabula Microcebus. **a**, **b**, **c**, UMAP plots of scRNA-seq datasets colored by compartment (**a**), by tissue (**b**) and by individual (**c**) after integration using FIRM, Seurat, Harmony, BBKNN and Scanorama.

We also evaluated FIRM’s ability to integrate large datasets across different technology platforms for the entire organism to construct a comprehensive atlas. To this end, we integrated 44,779 cells profiled using SS2 for all tissues with 54,865 cells profiled using 10X for all tissues in Tabula Muris (Fig. 6). For the shared cell populations across platforms, FIRM shows extensive mixing performance. The tissuespecific cells in that were found only in SS2 data remained correctly unmixed after integration: for example, microglial cells in brain myeloid; oligodendrocytes in brain non-myeloid; cells in large intestine; keratinocyte stem cells in skin.

**Fig. 6.**
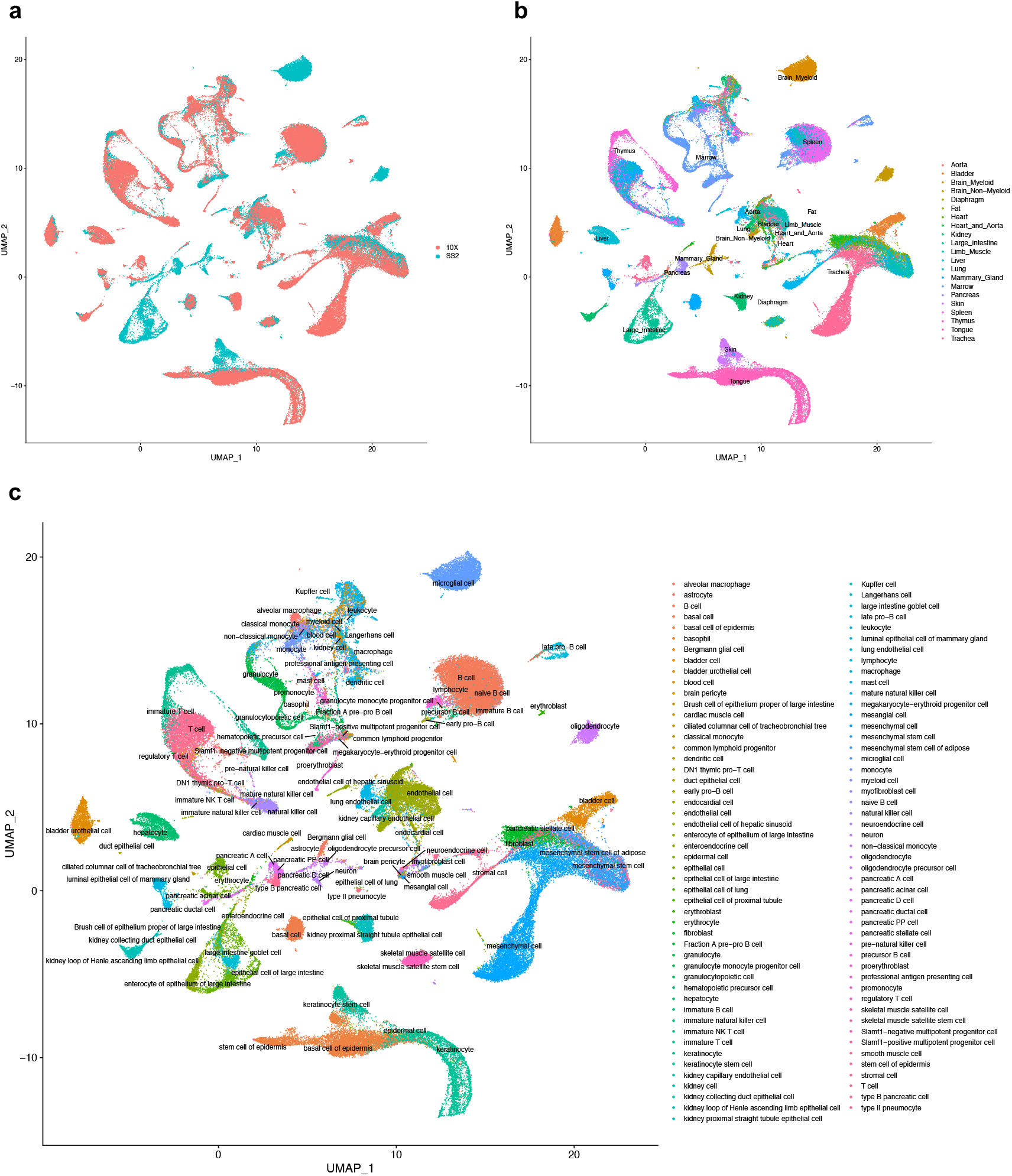
The performance of FIRM for integrating the whole SS2 dataset and 10X dataset of the entire organism in Tabula Muris. **a**, **b**, **c**, UMAP plots of scRNA-seq datasets colored by platform (**a**), by tissue (**b**) and by cell type (**c**) after integration using FIRM.

The FIRM algorithm naturally allows parallelization, because it gradually changes the number of clusters in SS2 and 10X datasets and searches the combination that gives the best cluster alignment. Therefore, the computational time of FIRM can be greatly shortened by using more CPU cores. The time taken to integrate 44,779 cells profiled using SS2 with 54,865 cells profiled using 10X for all tissues in Tabula Muris using FIRM took about 1-3 hours, varying based on the number of clusters in the 10X dataset. For integration of 12,329 SS2 cells and 231,752 10X cells in Tabula Microcebus, FIRM took about 5-8 hours. The overall computational time also depends on the total number of cells. We evaluated the computational time for the SS2-10X integration of Tabula Muris using 30 CPU cores. The time varies from one minute to two hours for different tissues with number of cells ranging from 934 to 12,598 (Supplementary Fig. 34; SI embargoed until publication).

## Discussion

FIRM is an accurate and effective method for integrating scRNA-seq datasets across multiple tissue types, experiments, and platforms. The integrated dataset can help to answer relevant biological questions and increase the confidence of analytical conclusions. For downstream analysis to be biologically meaningful, it is important to minimize technical variations such as batch effects while preserving biological variations of interest. Generally, it is very difficult to distinguish technical from biological variation, and overcorrection can occur when attempting to remove technical variation, resulting in loss of critical underlying biological variations. The best way to avoid overcorrection is to design methods that target minimization of specific types of confounding variation. FIRM successfully does so by specifically accounting for the heterogeneity in cell type composition between datasets which is a hurdle in efficient data integration. FIRM not only adjusts for the effect of cell type composition but also preserve the biological differences, therefore FIRM can be applied to integrate datasets across different individuals and tissues to study the biological differences across samples and tissues, in addition to datasets across different platforms. Other existing integration methods that use a general approach to account for variation between datasets do so by aligning cells with high similarity, and as such they are prone to inadvertently removing the biological differences across individuals as well. In contrast with existing methods, FIRM requires no assumption about shared cell populations between datasets, and is therefore applicable even without prior knowledge about the dataset composition.

FIRM integrates datasets based on the intersection of highly variable genes in each dataset. This preserves the most important expression patterns with less noise for alignment. However, we cannot ensure that the expressions of all genes are harmonized between SS2 cells and 10X cells, especially for low abundance transcripts. This is a remaining challenge to be addressed by future method development.

Through analysis of a diverse collection of human, mouse, and mouse lemur datasets, we show that FIRM outperforms or performs comparably to existing methods in terms of accuracy of integration and superior preservation of original structure for each dataset. Ultimately, our data integration tool enables new biological insights, and provides efficiency and utility for large scale projects.

## Methods

### Data preprocessing

For all scRNA-seq datasets, we performed the standard pre-processing workflow in Seurat^21^, which includes normalization, scaling and feature selection.

More specifically, for each dataset, we employed the log-normalization which is the default normalization method in Seurat. We used the gene expression matrix ***X***, where *X_ij_* is the number of reads (for SS2) or unique molecular identified (UMI, for 10X) for gene *i* that are detected in cell *j*. For each cell, the feature counts are divided by the total counts for that cell, multiplied by a scale factor *M* and then transformed using 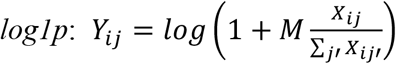.

Then we scaled the expression values for each gene across all cells in each dataset so that each gene has unit variance. Different from the default scaling method in Seurat, we did not center the expression to have zero mean across cells for the convenience of the re-scaling step in “Re-scaling via subsampling and generation of integrated data” that will follow.

In order to highlight the biological signal in scRNA-seq datasets, we identified genes with high variability across cells. For each dataset, we implemented the ‘FindVariableFeatures’ function in Seurat to select highly variable genes, where it ranked genes according to the dispersion after controlling the mean expression. In default, we selected the top 4,000 genes. For integrative analysis of two datasets across platforms, we selected genes that are highly variable in both datasets.

### Cell clustering for each data set

As the analysis in this paper is unsupervised, where we did not use the information of cell type annotations when implementing the methods, we need to cluster cells for each dataset first and then we can align the cell clusters across datasets. Before cell clustering, we first performed dimensionality reduction to obtain the low-dimensional embedding for each cell. Specifically, we performed PCA for each dataset, where the scaled data with the highly variable genes is used. We chose the number of PCs according to its relationship with the variance explained. For integrative analysis of two datasets across platforms, the number of PCs need to be the same. Typically, more PCs are needed for larger datasets. For the scRNA-seq datasets analyzed in this paper, we found that the performance of FIRM is robust to the number of PCs. We chose the number of PCs as the larger number in the original analyses that were performed separately on SS2 and 10X datasets^12,34^. Then for each dataset, we clustered cells based on their PC scores using the clustering approach in Seurat which identify clusters of cells by a shared nearest neighbor (SNN) modularity optimization-based clustering algorithm. The resolution parameter in the ‘FindClusters’ function in Seurat is used to control the number of clusters, which is tuned in FIRM for better integration.

### Cluster alignment via subsampling

After clustering the cells in 10X data and SS2 data separately, our next step is to align the cell clusters that represent the same cell types across the two datasets. We checked the alignment via subsampling to avoid overcorrection. First, we concatenated the scaled SS2 and 10X data and performed PCA on the combined data to obtain the low-dimensional representations for each cell. Next, we attempt to align each 10X cluster with a SS2 cluster in the following steps.

1. For 10X cluster *a* and the SS2 cluster *b,* where SS2 cluster *b* is among the five nearest SS2 clusters to 10X cluster *a*, we calculated the distance between their centers: 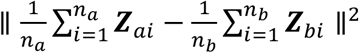. ***Z***_*a*_, ***Z***_*b*_ are the cells, and *n_a_*, *n_b_* are the numbers of cells in 10X cluster *a* and SS2 cluster *b*, respectively.
2. We then calculated the 75% quantile among all distances from the cells in SS2 cluster *b* to their cluster center: 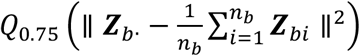. We check if the following criterion holds

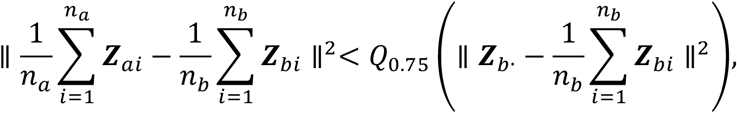
3. We considered the nearest SS2 cluster, among the five nearest SS2 clusters to 10X cluster *a*, that also satisfied the criterion in step (2) to be aligned with 10X cluster *a*.

However, even the same cell type may not be aligned in the previous procedure because of the difference in its abundances in 10X and SS2 data. To address this issue, for the 10X clusters which had not been aligned, we further performed subsampling to adjust the proportions of the 10X and SS2 clusters being considered and checked the alignment. For example, when we consider the 10X cluster *a* and the SS2 cluster *b*, if the proportion of 10X cluster *a* in 10X dataset is larger than the proportion of SS2 cluster *b* in SS2 dataset, i.e., 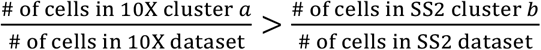, we subsampled the cells in 10X cluster *a* to obtain a subset, so that the proportion of 10X cluster *a* in this subset was the same with the proportion of SS2 cluster *b* in SS2 dataset, 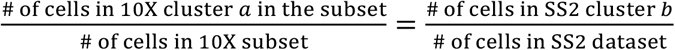. We then calculated the standard deviation of each gene across cells in this subset based on the original scaled expression values, i.e., *s_j_* = *sd*(*Z_ij_*), *i* ∈ 10X subset, where *Z_ij_* is the scaled data after the preprocessing procedure described in the “Data preprocessing” step. We performed re-scaling for cells in the whole 10X dataset using this standard deviation, i.e., 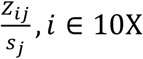 dataset. Based on the re-scaled data, we checked the alignment again using steps (1-3) described above. After the cluster alignment step, clusters of the cell types which existed in both 10X and SS2 datasets were expected to be aligned.

### Re-scaling via subsampling and generation of integrated data

To calculate the scaling factor for effective re-scaling, we need two datasets, one SS2 dataset and one 10X dataset, which contain the same types of cells and have the same cell-type proportions as well. Based on the aligned clusters identified using the above procedure, we performed subsampling for cells in the SS2 and 10X cluster pairs to obtain the SS2 subset and the 10X subset which meet our requirements. The subsampled datasets did not contain cells in the unaligned clusters, because unaligned clusters were considered as unique cell types which only existed in one dataset. Based on each of the subsampled datasets, we computed the standard deviations for each gene across cells on the original scaled expression values, i.e., *s*_*SS*2 *j*_ $ = *sd*(*Z_ij_*), *i* ∈ SS2 subset, and *s*_10X *j*_ $ = *sd*(*Z_ij_*), *i* ∈ 10X subset, where *Z_ij_* is the scaled data after the preprocessing procedure described in the “Data preprocessing” step. We used the calculated standard deviations to re-scale the gene expression values for cells in the whole SS2 and 10X datasets. i.e., 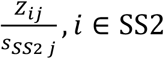 dataset and 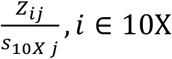 dataset. We concatenated the re-scaled data directly to obtain the integrated data.

### Integration Metrics

#### Mixing metric

We used the mixing metric designed in Seurat^21^ to evaluate how well the datasets mixed after integration. If the local neighborhood for a cell is well mixed across datasets, at least a small number (*k* = 5) of cells from each dataset is assumed to be its neighbors. For each cell, we obtain its (*k.max* = 300) ranked nearest neighbors in the integrated dataset and record the rank of the 5th nearest neighbor in each dataset (with a max of 300). The average of the ranks across all cells is defined as the mixing metric. As a result, smaller mixing metric typically indicates better mixing.

#### Local structure metric

We used the local structure metric designed in Seurat^21^ to determine how well the original structure of each dataset was reserved after integration. For each cell, we compare its *k* = 20 nearest neighbors in the original dataset and the integrated dataset. The average value of the fraction of overlapped neighbor across all cells is defined as the local structure metric. A large local structure metric indicates good preservation.

#### Average silhouette width (ASW)

The silhouette width for cell *i* from cell type *c* is 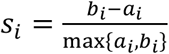 where *a_i_* is the average distance from cell *i* to all cells in cell type *c*, *b_i_* is lowest value of average distances from cell *i* to all cells for each cell type other than *c*. ASW is the mean of silhouette widths across all cells, where a higher score indicates cells are closer to cells of the same cell type and are further from cells of different cell types. We calculated ASW based on the predefined cell identities and low-dimensional embedding space of the integrated dataset using PCA.

#### Adjusted rand index (ARI)

ARI measures the similarity between two clustering results. The ARI is defined as 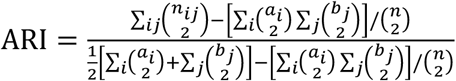 where *n_ij_*, *a_i_*, *b_j_* are values from the contingency table. For the integrated dataset, we clustered cells based on their PCA scores using the clustering approach in Seurat with default settings. Then we calculated ARI to compare the clustering of integrated data with the predefined cell types, where higher values indicate higher similarities.

#### Resolution in clustering

Cluster alignment is the key for effective integration. The resolution parameter in the ‘FindClusters’ function in Seurat is used to control the number of clusters. Unsuitable resolutions for clustering of SS2 and 10X datasets may lead to bad integration results. We searched the best resolution pairs for SS2 and 10X datasets in a default range of 0.1 to 2. Under each resolution pair, we integrated the datasets using our method and calculated the mixing metric. As our method would not suffer overcorrection, smaller mixing metric always indicates better integration. Therefore, we chose the pair yielding the smallest mixing metric.

#### Baseline model

We considered the special case without the re-scaling procedure to be the baseline model. We directly concatenated the scaled expression matrix for the overlapped highly variable genes after data processing to obtain the integrated dataset. If the mixing metric of integrated dataset after re-scaling does not decrease, we chose the baseline model.

#### Label transfer and match scores

The integration of datasets enables efficient label transfer between datasets. Suppose we want to use the annotations for cells in 10X dataset to annotate cells in SS2 dataset. For each SS2 cell, we found its 10 nearest 10X cells in the integrated dataset and summarized the cell types they belong to. We chose the cell type with the highest frequency to annotate the SS2 cell.

In case that some cell types do not exist in 10X dataset, we defined the match score to measure whether the cell in SS2 is present in 10X data. For each SS2 cell, we divided its averaged distance to its 10 nearest neighbors in SS2 dataset by that in 10X dataset. Lower score means less likely to be present in 10X data.

#### Datasets

Below we describe all of the datasets used in the current work.

#### Tabula Muris

The Tabula Muris dataset^12^ contains 44,949 cells profiled using SS2 from 20 organs and 55,656 cells profiled using 10X from 12 mouse organs. We removed cells without annotations in the original study^12^. The filtered dataset contained 44,779 cells profiled using SS2 and 54,865 cells profiled using 10X.

#### Tabula Microcebus

The Tabula Microcebus dataset contains 12,329 cells profiled using SS2 from 20 organs and tissues in 3 individuals and 231,752 cells profiled using 10X from 25 organs and tissues in 4 individuals which were annotated in the original study.

#### Human Lung Atlas

We considered the scRNA-seq data of Patient 1 in the Human Lung Atlas dataset^34^, which contains 3,987 cells profiled using SS2 and 9,744 cells profiled using 10X which were annotated in the original study^34^.

## Data availability

The datasets in Tabula Muris used in this manuscript are available at http://tabulamuris.ds.czbiohub.org/. The Human Lung Atlas data is available on Synapse (https://www.synapse.org/#!Synapse:syn21041850). The datasets in Tabula Microcebus are from the Tabula Microcebus consortium.

## Code availability

FIRM code is available as Supplementary Code and at https://github.com/mingjingsi/FIRM.

## Acknowledgements

This work is supported in part by Hong Kong Research Grant Council [16101118, T12-704/16R-2, 24301419, 14301120], the Hong Kong University of Science and Technology’s startup grant [R9364], the Hong Kong University of Science and Technology Big Data for Bio Intelligence Laboratory (BDBI), the Lo Ka Chung Foundation through the Hong Kong Epigenomics Project, the Chau Hoi Shuen Foundation, the ASPIRE League through a Seed Grant Award, the Chinese University of Hong Kong direct grants [4053360, 4053423], the Chinese University of Hong Kong startup grant [4930181], the Chinese University of Hong Kong’s Project Impact Enhancement Fund (PIEF) and Science Faculty’s Collaborative Research Impact Matching Scheme (CRIMS), the East China Normal University startup grant, the Shanghai Sailing Program.

